# Proline provides a nitrogen source in the retinal pigment epithelium to synthesize and export amino acids for the neural retina

**DOI:** 10.1101/2023.04.18.537355

**Authors:** Siyan Zhu, Rong Xu, Abbi L. Engel, Yekai Wang, Rachel McNeel, James B. Hurley, Jennifer R. Chao, Jianhai Du

**Author notes:** Corresponding Authors: Jianhai Du, One Medical Center Dr, PO Box 9193, WVU Eye Institute, Morgantown, WV 26505. These authors contributed equally to this work (Siyan Zhu, Rong Xu).

## Abstract

It is known that metabolic defects in the retinal pigment epithelium (RPE) can cause degeneration of its neighboring photoreceptors in the retina, leading to retinal degenerative diseases such as age-related macular degeneration. However, how RPE metabolism supports the health of the neural retina remains unclear. The retina requires exogenous nitrogen sources for protein synthesis, neurotransmission, and energy metabolism. Using ^15^N tracing coupled with mass spectrometry, we found human RPE can utilize the nitrogen in proline to produce and export 13 amino acids, including glutamate, aspartate, glutamine, alanine and serine. Similarly, we found this proline nitrogen utilization in the mouse RPE/Cho but not in the neural retina of explant cultures. Co-culture of human RPE with the retina showed that the retina can take up the amino acids, especially glutamate, aspartate and glutamine, generated from proline nitrogen in the RPE. Intravenous delivery of ^15^N proline *in vivo* demonstrated ^15^N-derived amino acids appear earlier in the RPE before the retina. We also found proline dehydrogenase (PRODH), the key enzyme in proline catabolism, is highly enriched in the RPE but not the retina. The deletion of PRODH blocks proline nitrogen utilization in RPE and the import of proline nitrogen-derived amino acids in the retina. Our findings highlight the importance of RPE metabolism in supporting nitrogen sources for the retina, providing insight into understanding the mechanisms of the retinal metabolic ecosystem and RPE-initiated retinal degenerative diseases.

## Introduction

The retinal pigment epithelium (RPE) is essential for maintaining metabolic homeostasis of the outer retina. The RPE relies on an active metabolism to transport nutrients from the choroidal blood circulation to the metabolically demanding photoreceptors. It also transports “metabolic waste” from the photoreceptors to the blood, phagocytoses daily-shed photoreceptor tips, and synthesizes ketone bodies and mitochondrial intermediates for photoreceptor utilization. Defects in RPE metabolism could lead to disruption of the metabolic ecosystem in the outer retina, contributing to multiple types of retinal degenerative diseases including age-related macular degeneration, the leading cause of blindness in the elderly (1-3).

The RPE is capable of using multiple types of nutrient sources to fuel its metabolism which spares glucose for the neural retina (4). Our previous research demonstrates that the RPE prefers to use proline, a non-essential amino acid as a metabolic substrate (5). RNA-Sequence data also shows that genes in proline transport and catabolism are highly enriched in human RPE (6). Proline supplementation decreases the effects of oxidative damage in cultured RPE, and dietary proline protects against retinal degeneration induced by oxidative damage of RPE *in vivo* (7). Aside from being provided by the diet, proline can be produced from ornithine, glutamate or collagen degradation. Mutations of enzymes in these pathways are known to cause RPE cell death and retinal degeneration (8). For example, mutations of ornithine aminotransferase (OAT) cause gyrate atrophy, an ocular disease characterized by progressive loss of the RPE/Cho, leading to retinal degeneration. Proline is catabolized through proline dehydrogenase (PRODH) in the mitochondrial matrix into pyrroline-5-carboxylate (P5C), which is further degraded into glutamate, a substrate for the tricarboxylic acid (TCA) cycle. By ^13^C tracing, we have found the carbon skeleton from proline is catabolized into glutamate and TCA cycle intermediates which are exported from the retinal side of the RPE to support retinal metabolism (7). However, the fate of the nitrogen groups in proline catabolism remains unclear.

In addition to high metabolic needs for carbon sources such as glucose, the neural retina also requires nitrogen sources to produce ATP, cGMP, glutamate and other nitrogen-containing metabolites for energy metabolism, phototransduction and protein synthesis. Our previous studies in both human and mouse explant cultures showed that retinas need substantial amounts of exogenous glutamate and aspartate (6). Glutamate and aspartate are interconvertible through aspartate transaminase. Glutamate is a key neurotransmitter as well as a metabolite substrate for the TCA cycle and many amino acids such as serine, glycine, glutamine and γ-aminobutyric acid (GABA). The retinal neurons contain millimolar concentrations of glutamate. However, glutamate and aspartate are the least abundant amino acids in plasma, suggesting they might be produced locally in the neural retina or the RPE.

In this report, we trace the fate of nitrogen from proline in the RPE and the neural retina by using ^15^N proline. We found that proline is a nitrogen source for synthesizing 13 amino acids, especially glutamate, aspartate and glutamine in primary human RPE. However, the neural retina alone cannot utilize the nitrogen directly from proline. Interestingly, neural retina co-cultured with RPE imports amino acids that were made by the RPE using nitrogen from proline. Additionally, we found the neural retina can utilize nitrogen from proline *in vivo* to produce glutamate, aspartate and other amino acids by proline catabolism through PRODH in the RPE. Our results demonstrate that proline catabolism in the RPE is a key source of nitrogen for the synthesis of important amino acids for the neural retina.

## Results

### Proline is an important nitrogen source for the synthesis and export of amino acids in human RPE cells

To trace the fate of nitrogen in proline catabolism, we supplemented mature primary human RPE cells in culture with or without ^15^N proline. The spent media and cells were harvested to analyze ^15^N labeled amino acids using gas chromatography mass spectrometry (GC MS) (**Fig 1A**). The nitrogen from proline can be utilized to synthesize 13 types of amino acids (9) (**Fig 1B**). Therefore, we targeted these amino acids together with proline in our GC MS analysis. As expected, human RPE cells actively consumed proline and 1mM ^15^N proline was almost completely used up after incubation for 48h (**Fig S1A**). ^15^N proline was incorporated in all the selected amino acids including glutamine, aspartate, glutamate, alanine, oxoproline, branched-chain amino acids (BCAAs, leucine, valine and isoleucine), ornithine, asparagine, γ-aminobutyric acid (GABA), serine and glycine (**Fig 1C**). Glutamine and aspartate had the highest enrichment, whereas glycine had the lowest. Proline also increased the cellular pool sizes of some amino acids, especially aspartate and asparagine (**Fig 1D**). The RPE is known to actively transport nutrients to support the neural retina. To investigate whether the RPE can export these proline-derived amino acids out of the cells, we quantified these amino acids in the spent media. Except for GABA, all the other amino acids were labeled by ^15^N proline in the spend media at 24h, and the enrichment was further increased at 48h (**Fig 1E**). Similar to the intracellular changes, glutamine, aspartate, alanine, oxoproline, glutamate and BCAAs were highly enriched by ^15^N proline in the media. The total amounts of these amino acids were only slightly increased or unchanged except for aspartate, which increased ∼4-6 fold by ^15^N proline (**Fig 1F**). These results suggest that proline provides an important nitrogen source for the synthesis of multiple amino acids, which might be exported to support retinal metabolism.

**Figure 1.**
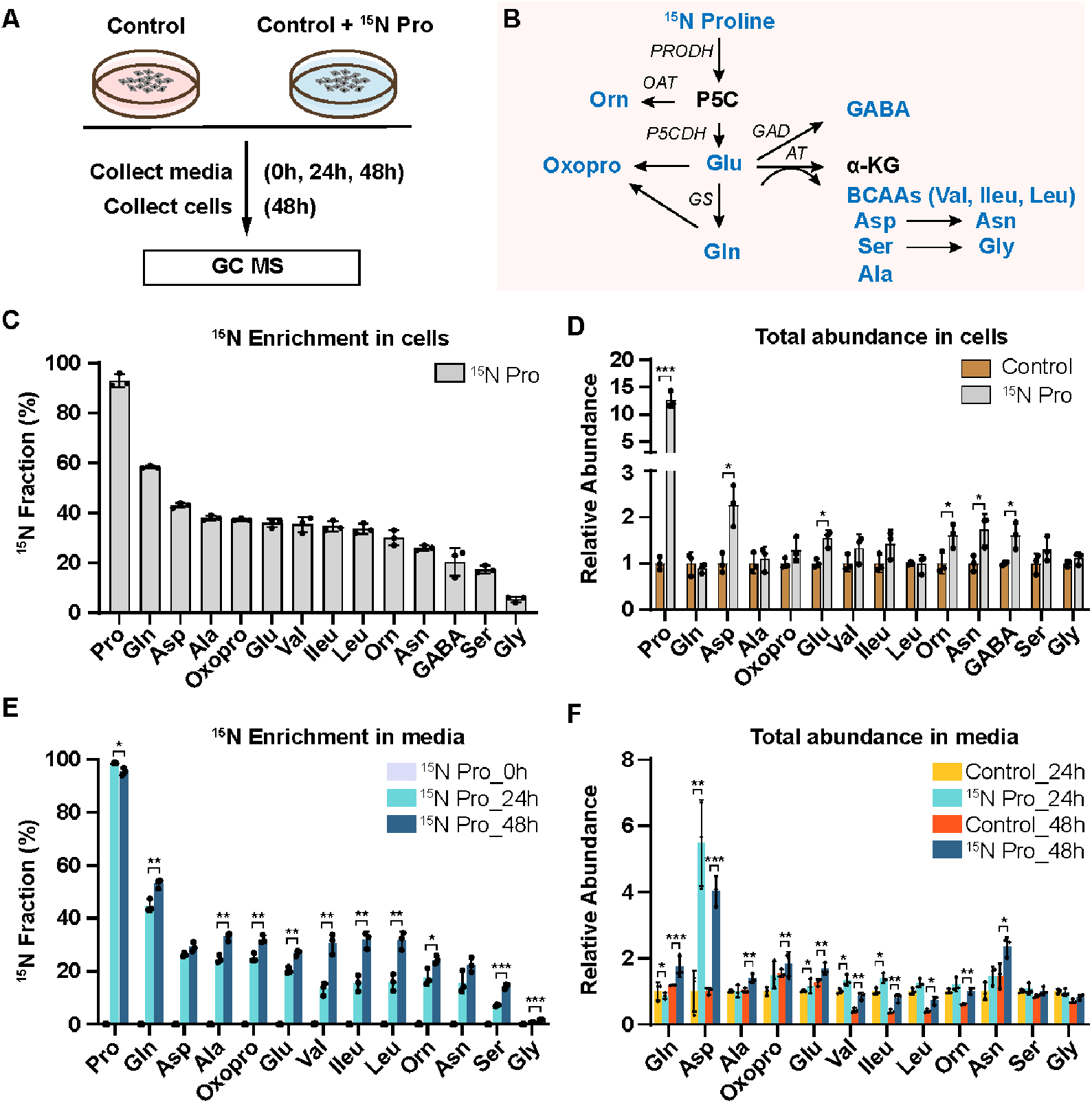
Human RPE utilizes nitrogen from proline to generate amino acids and exports them into the media. (A) Human RPE cells were grown for 20 weeks and switched into DMEM media with or without 1 mM ^15^N proline. The spent media were collected at 0h, 24h and 48h and cells were collected at 48h for metabolite analysis with GC MS. (B) A schematic of proline nitrogen metabolism into amino acids (Colored in blue) through proline dehydrogenase (PRODH), pyrroline-5-carboxylic acid (P5C) dehydrogenase (P5CDH), ornithine transaminase (OAT), glutamate decarboxylase (GAD), glutamine synthetase (GS) and aminotransferases (AT). These amino acids in proline metabolism were selected for GC MS analysis. (C-D) ^15^N fraction of metabolites derived from ^15^N proline in RPE cells and the total abundance of all isotopologue relative to control RPE cells without ^15^N proline. *P<0.05, ***P<0.001, N=3. (E-F) ^15^N fraction of metabolites derived from ^15^N proline in RPE spent media at different time points and the total abundance of all isotopologue relative to control RPE culture media without ^15^N proline at 24h or 48h. *P<0.05, **P<0.01, ***P<0.001, N=3.

### RPE, but not the neural retina, is responsible for the amino acid synthesis by proline

We previously reported that proline is primarily catabolized in the RPE to produce TCA cycle intermediates, but proline catabolism in the neural retina is negligible (7). To investigate the sites of amino acid synthesis from the nitrogen group in proline, we incubated freshly isolated mouse RPE/Cho and the neural retina with ^15^N proline in the presence of 5 mM glucose in Krebs-Ringer Bicarbonate (KRB) buffer. Then we collected the tissues within 2 hours to analyze ^15^N proline-derived amino acids (**Fig 2A**). ^15^N proline in the media was consumed 4-5 times faster in the RPE/Cho than the neural retina (**Fig 2B**). Consistent with previous findings, ^15^N from the labeled proline was incorporated rapidly into 9 types of amino acids in the RPE/Cho. Similar to human RPE, glutamate, aspartate, glutamine, oxoproline and alanine were highly enriched with ^15^N from proline (**Fig 2C-J**). The total abundance of these amino acids remained constant during the 2h incubation, except for the slight increase of aspartate and decrease of leucine (**Fig S2A**). These results further support the previous findings that proline provides important nitrogen sources for amino acid synthesis in the RPE. By contrast, except for a very slight enrichment in alanine, we did not detect enrichment in the other amino acids in the neural retina (**Fig 2C-J**), demonstrating that the neural retina is incapable of utilizing proline as a nitrogen source directly. Proline was not sufficient to maintain the total concentrations of amino acids in the neural retina. In contrast to the RPE/Cho, the total abundance of several amino acids was diminished 20-60% after 2 hours of incubation (**Fig S2B**). These results suggest that while the neural retina requires exogenous amino acids other than proline to support its metabolism, it does not directly utilize proline.

**Figure 2.**
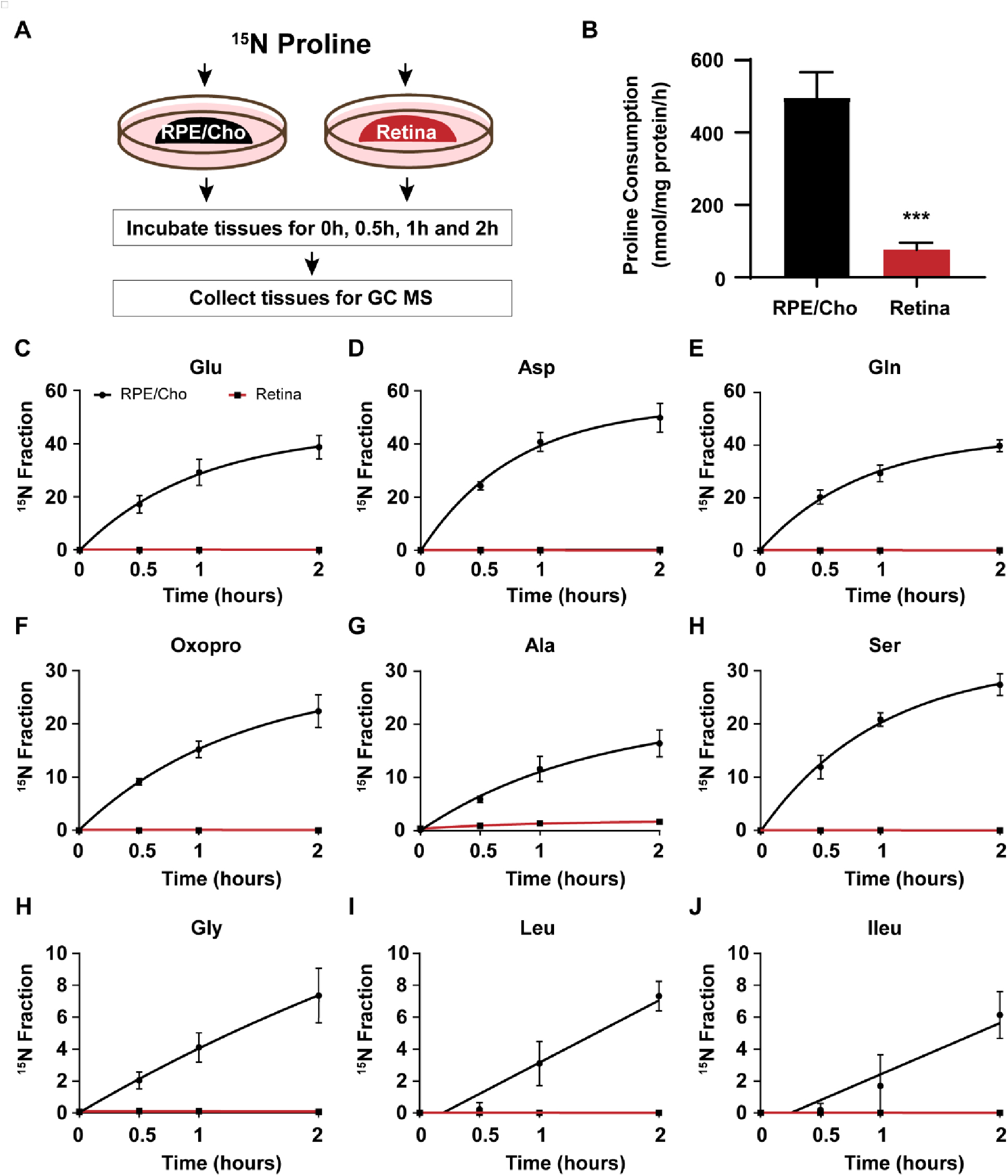
The synthesis of amino acids by proline occurs mostly in the RPE but not the neural retina ex vivo in mice. (A) A schematic of *ex vivo* incubation of mouse RPE/ Choroid (RPE/Cho) and retina with ^15^N proline. Tissues were incubated with ^15^N proline in Krebs-Ringer Bicarbonate Buffer (KRB) for 0h, 0.5h, 1h and 2h. Tissues and media were collected at these time points for metabolite analysis with GC MS. (B) ^15^N Proline consumption from RPE/Cho and retina. ***P<0.001, N=4. (C-J) ^15^N fraction of metabolites derived from ^15^N proline in RPE/Cho and retina at different time points. A) A schematic of ex vivo incubation of mouse RPE/Cho and retina with ^15^N proline. Both tissues were incubated with 1 mM ^15^N proline in KRB containing 5 mM glucose for 0h, 0.5h, 1h and 2h. Tissues and media were collected at these time points for metabolite analysis with GC MS. (B) ^15^N Proline consumption of RPE/Cho and retina from the media. ***P<0.001, N=4. (C-J). ^15^N fraction of metabolites derived from ^15^N proline in RPE/Cho and retina at different time points, N=4.

### Proline nitrogen-derived amino acids from the RPE can be utilized by the neural retina in co-culture

To test whether proline nitrogen-derived amino acids produced from the RPE can be utilized by the neural retina, we co-cultured human RPE cells with mouse retina for 6 hours in a medium supplemented with ^15^N proline and cultured human RPE or mouse retina alone under the same conditions as the control groups. The spent medium, RPE cells and retinas were harvested to analyze proline-derived amino acids by GC MS. (**Fig 3A**). As expected, except for ^15^N proline itself, the neural retina culture alone could not incorporate the ^15^N from proline into other amino acids. However, co-cultured retina had substantial ^15^N enrichment in multiple amino acids including glutamine, aspartate, alanine, glutamate, asparagine valine, serine and glycine (**Fig 3B**). Consistently, compared with RPE culture alone, most of these amino acids showed decreased ^15^N enrichment in co-cultured RPE. Compared to the retina or RPE culture alone, the co-culturing substantially increased the total abundance (concentrations) of almost all the amino acids except GABA in the retina but reduced these amino acids in the RPE (**Fig 3C**). These results suggest the retina imports these amino acids produced by the RPE. Similarly, RPE culture alone exported proline-derived amino acids into the media; however, many of these amino acids were reduced in the co-cultured media. (**Fig 3D-E**). Taken together, we confirm that the neural retina cannot catabolize proline directly, but it can import proline-derived amino acids from the RPE. (**Fig 3F**).

**Figure 3.**
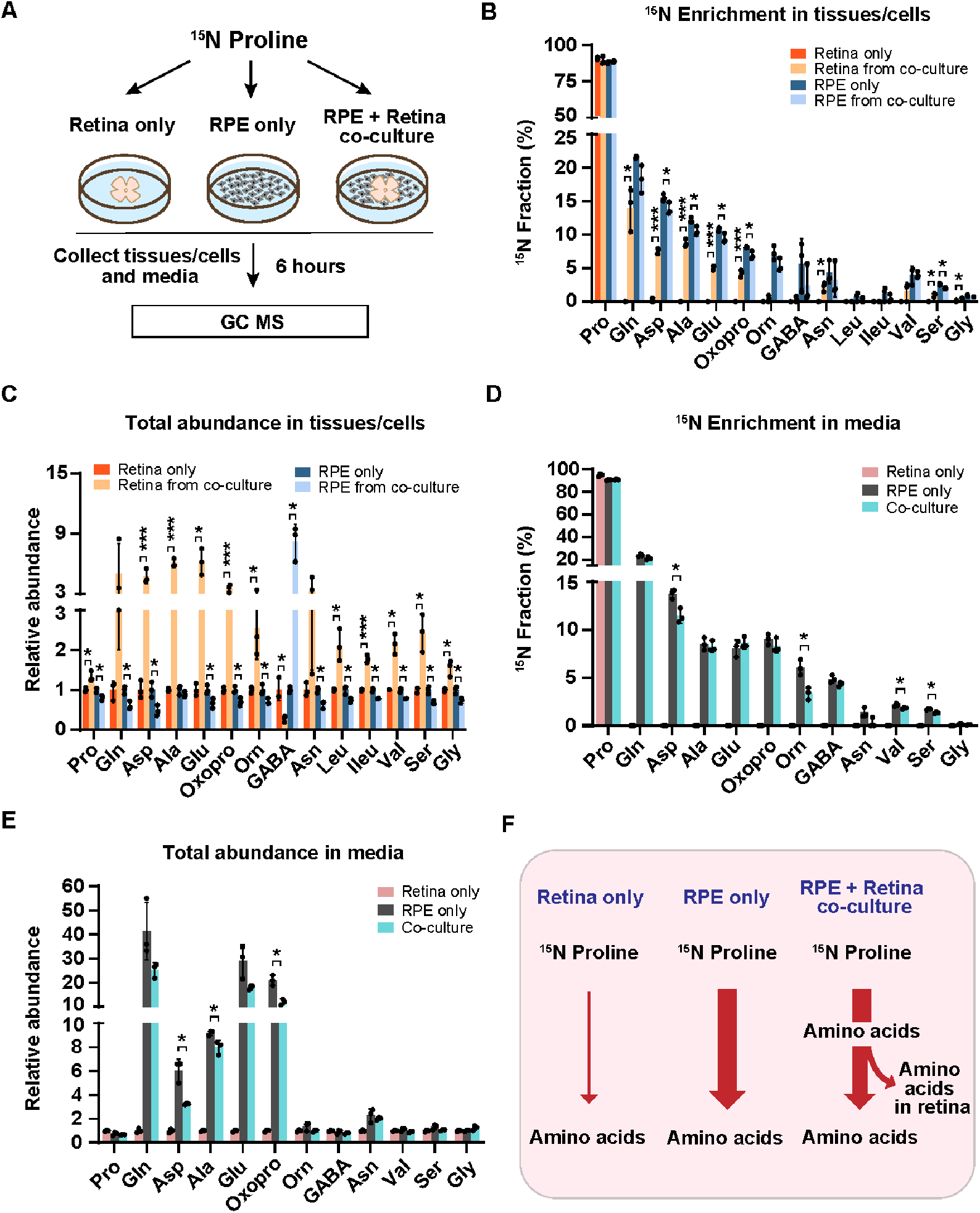
The neural retina imports proline-derived amino acids from the RPE. (A) Human RPE cells and mouse retina were cultured separately or co-cultured in DMEM media with 1mM ^15^N proline for 6 hours. Tissues and media were collected for metabolite analysis with GC MS. (B-C) ^15^N fraction of metabolites derived from ^15^N proline in RPE cells/retina and the total abundance of all isotopologue in RPE cells/retina from co-culture relative to their corresponding control. *P<0.05, ***P<0.001, N=3. (D-E) ^15^N fraction of metabolites derived from ^15^N proline in media and the total abundance of all isotopologue relative to control media from the retina-only group. *P<0.05, N=3. (F) A schematic of proline nitrogen metabolism in co-culture. Retina alone barely utilizes proline for amino acid synthesis, but RPE utilizes proline to produce amino acids to feed the retina.

### RPE exports proline-derived amino acids to the retina in vivo

To test whether the retina utilizes proline-derived amino acids from the RPE *in vivo*, we administered a bolus injection of ^15^N proline intravenously and analyzed ^15^N-proline-derived amino acids in the retina and RPE/Cho (**Fig 4A**). Except for ^15^N in proline, we detected only 6 amino acids (glutamine, oxoproline, aspartate, glutamate, alanine and serine) that were labeled by ^15^N proline, probably due to the rapid proline catabolism. The enrichment of ^15^N proline peaked at 5 min after injection in both the retina and RPE/Cho. However, the ^15^N enrichment of other amino acids peaked at 15 min in the RPE/Cho and 30 min in the neural retina, indicating sequential proline catabolism into amino acids from the RPE/Cho to the retina (**Fig 4C-H**). The ^15^N labeling of all these amino acids in RPE/Cho was higher and earlier than the retina. The RPE/Cho incorporated ^15^N into amino acid at 5 min, peaked at 15 min, and dropped at 30 min; however, in the retina, there was almost no ^15^N labeling at 5 min although the labeling appeared at 15 min, steadily increased, and peaked at 30 min. This data strongly suggests the RPE metabolizes proline first before exporting to the retina. Except for transient increases in proline, glutamine and aspartate, the concentrations of these amino acids were constant in the RPE/Cho and the retina at different time points after injection (**Fig S3A-B**). As the bolus injection had a lower rate of enrichment and labeled fewer amino acids, we administered a second retro-orbital injection of the same dose of ^15^N proline 10 min after the first injection and extended the collection time to 60 min (**Fig 5A**). Indeed, compared to the bolus injection alone, the double injection increased the number of ^15^N labeled amino acids from 6 to 10 and almost doubled the ^15^N enrichment (**Fig 5B-K**). Similar to the bolus injection, the labeled amino acids appeared earlier with higher enrichment in the RPE/Cho than the retina before 30 min. At 60 min, all the ^15^N-labeled amino acids, except for GABA, were decreased in the RPE/Cho, but continued to rise in the retina (**Fig 5B-L**). These time lags between the RPE and retina further suggest that proline is primarily catabolized in the RPE, and its derived amino acids are utilized by the retina. Compared to single injection, double injections have a more transient increase in the concentrations of proline-derived amino acids in the retina (**Fig S4A-B**), suggesting that proline is an important source of synthesis of these amino acids.

**Figure 4.**
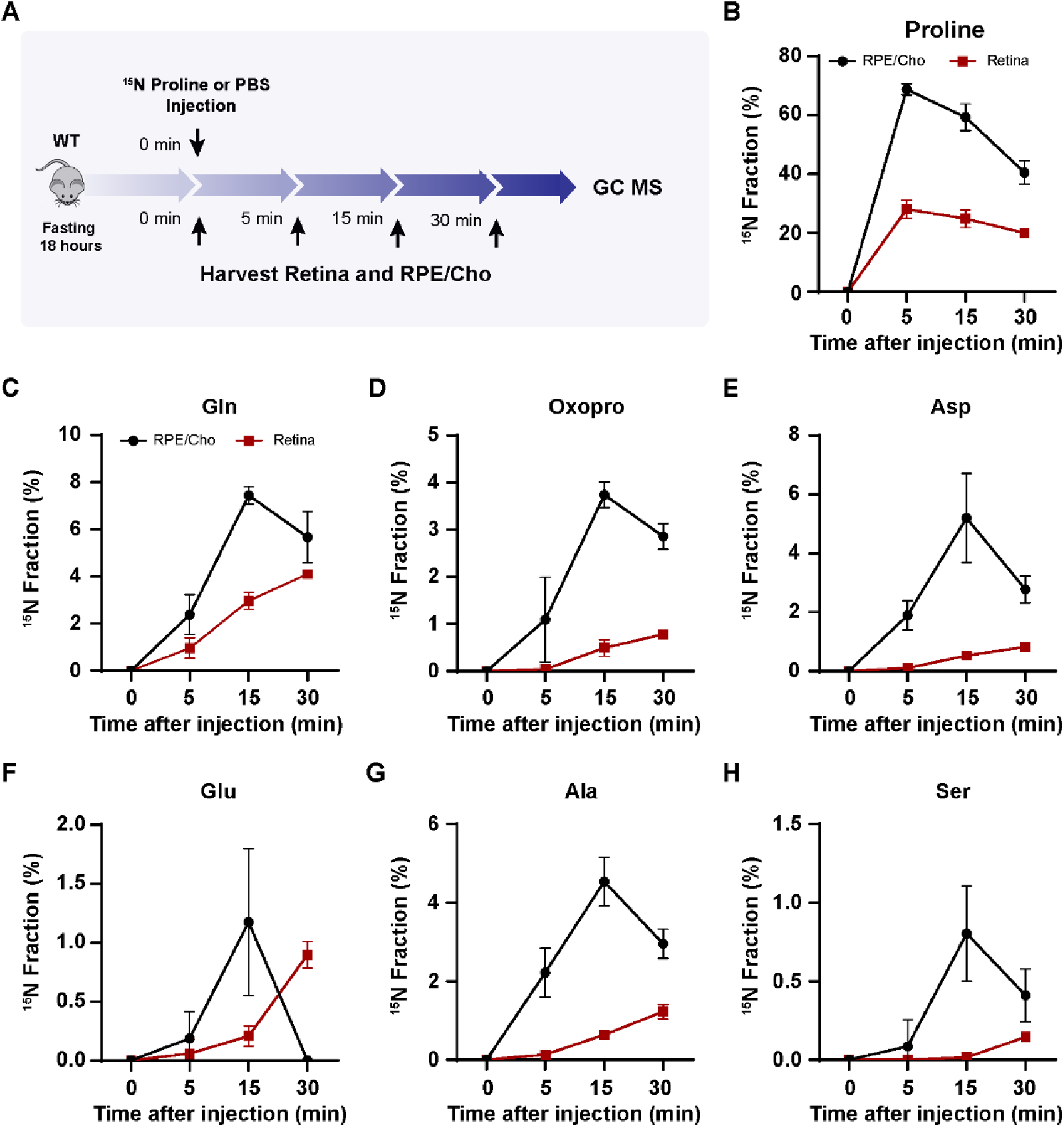
Proline-derived amino acids appear later in the retina than the RPE/Cho. (A) A schematic of *in vivo* tracing of ^15^N proline. Mice were fasted for 18 hours and received a single injection of ^15^N proline (150 mg/kg) or PBS. Retina and RPE/Cho were harvested at 0min, 5min, 15min and 30min after injection for metabolite analysis with GC MS. (B-H) ^15^N fraction of proline and metabolites derived from ^15^N proline in RPE/Cho and retina at different time points. N=4.

**Figure 5.**
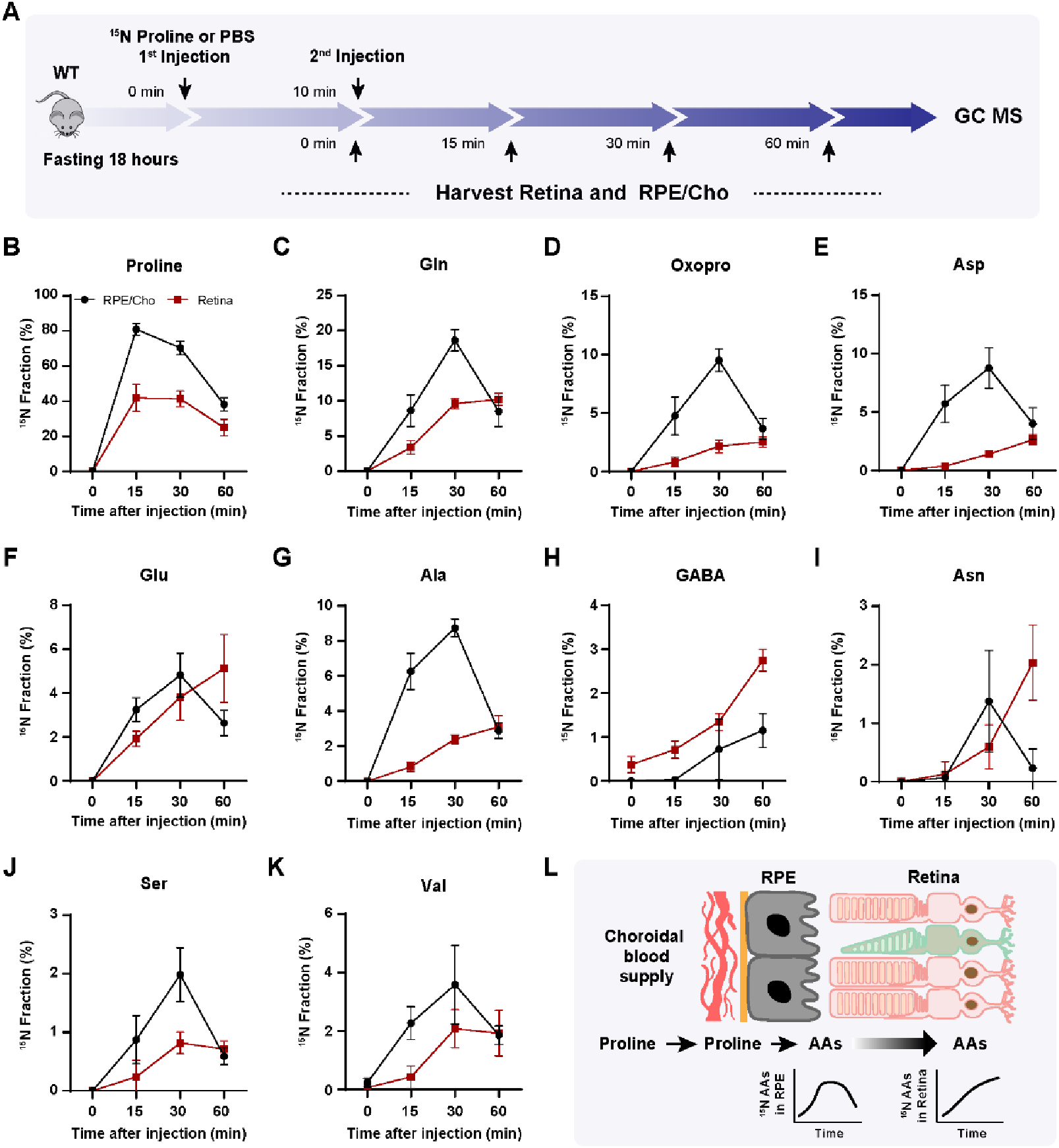
There is a delayed synthesis of amino acids from ^15^N proline in the retina compared to the RPE/Cho. (A) A schematic of *in vivo* tracing of ^15^N proline with double injection. Mice were fasted for 18 hours and received two injections of ^15^N proline (150 mg/kg) or PBS at 0 min and 10 min, respectively. Retina and RPE/Cho were harvested at 0min, 5min, 15min and 30min after the second injection for metabolite analysis with GC MS. (B-K) ^15^N fraction of proline and metabolites derived from ^15^N proline in RPE/Cho and retina at different time points. N=8 (L) A schematic of proline catabolism into amino acids in RPE to be exported into the neural retina.

### Loss of Prodh blocks the utilization of proline-derived amino acids in the retina from the RPE/Cho

To investigate whether proline catabolism in the RPE is critical for the amino acid supply in the retina, we tested the protein expression of PRODH, the key enzyme in proline catabolism. Immunoblotting showed that PRODH protein expression in RPE/Cho was 15 to 20-fold higher than the neural retina (**Fig 6A-B**), which is consistent with our previous results that ^15^N proline is primarily catabolized in the RPE/Cho but not the retina. The inbred mouse line PRO/ReJ (*Prodh*-/-) carries a G -> T substitution 135bp upstream of the native terminal codon in the *PRODH* gene, which leads to premature translational termination (10). Due to the lack of PRODH activity, this mouse line has hyperprolinemia (11). We validated that the PRODH protein expression was not detectable in the liver, retina and RPE/Cho of *Prodh*-/-mice (**Fig 6C-D**). Consistently, loss of PRODH caused substantial accumulation of proline in the liver, plasma, retina and RPE/Cho (**Fig 6E-H**). To study whether deletion of PRODH can block the synthesis and export of proline-derived amino acids from the RPE to the retina, we administered a double injection of ^15^N proline as mentioned before and collected the retina and RPE/Cho 30 min after the first injection (**Fig 7A-B**). Similarly, we detected 9 ^15^N-labeled amino acids in both the retina and RPE/Cho in the wild type (WT) mice with ^15^N enrichment ranging from 1-15%. RPE/Cho had higher enrichment of all the amino acids when compared to the retina. However, in the *Prodh-/-* mice, the ^15^N enrichment of these amino acids was significantly diminished to less than 1% (**Fig 7C-K**), demonstrating that proline catabolism in the RPE is required for the utilization of proline-derived amino acids in the retina. Compared to the WT mice, the *Prodh*-/-mice had lower concentrations of glutamate, alanine and aspartate in the RPE/Cho (**Fig S5A**), suggesting that proline is important to maintain the availability of these amino acids. Alanine also was lower in the retina of the *Prodh*-/-mice. However, several amino acids including glutamine, oxoproline and asparagine were increased in *Prodh*-/-retina (**Fig S5B**), indicating there might be compensatory pathways for the synthesis of these amino acids.

**Figure 6.**
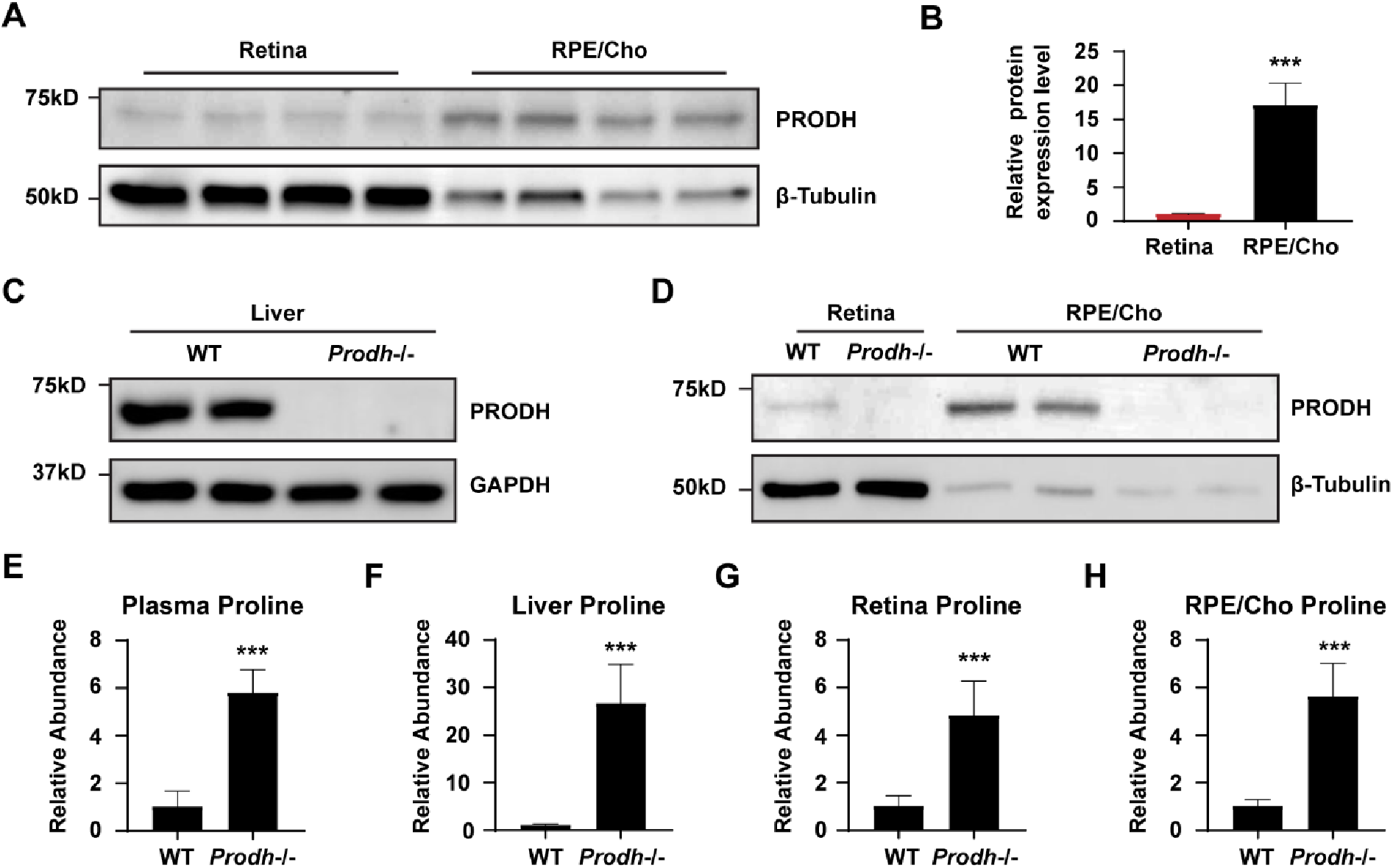
PRODH is more abundant in the RPE/Cho and deletion of *Prodh* causes proline accumulation. (A) RPE/Cho has higher PRODH protein expression than RPE/Cho. β-tubulin is used as the loading control. (B) The relative protein expression level of PRODH in retina and RPE/Cho. ***P<0.001, N=4. (C-D) *Prodh* mutation causes the depletion of PRODH proteins in liver, retina and RPE/Cho in *Prodh*-/-mice. (E-H) Total abundance of proline in the liver, plasma, PRE/Cho and retina from *Prodh*-/-mice relative to those tissues from WT mice. ***P<0.001, N≥4.

**Fig. 7.**
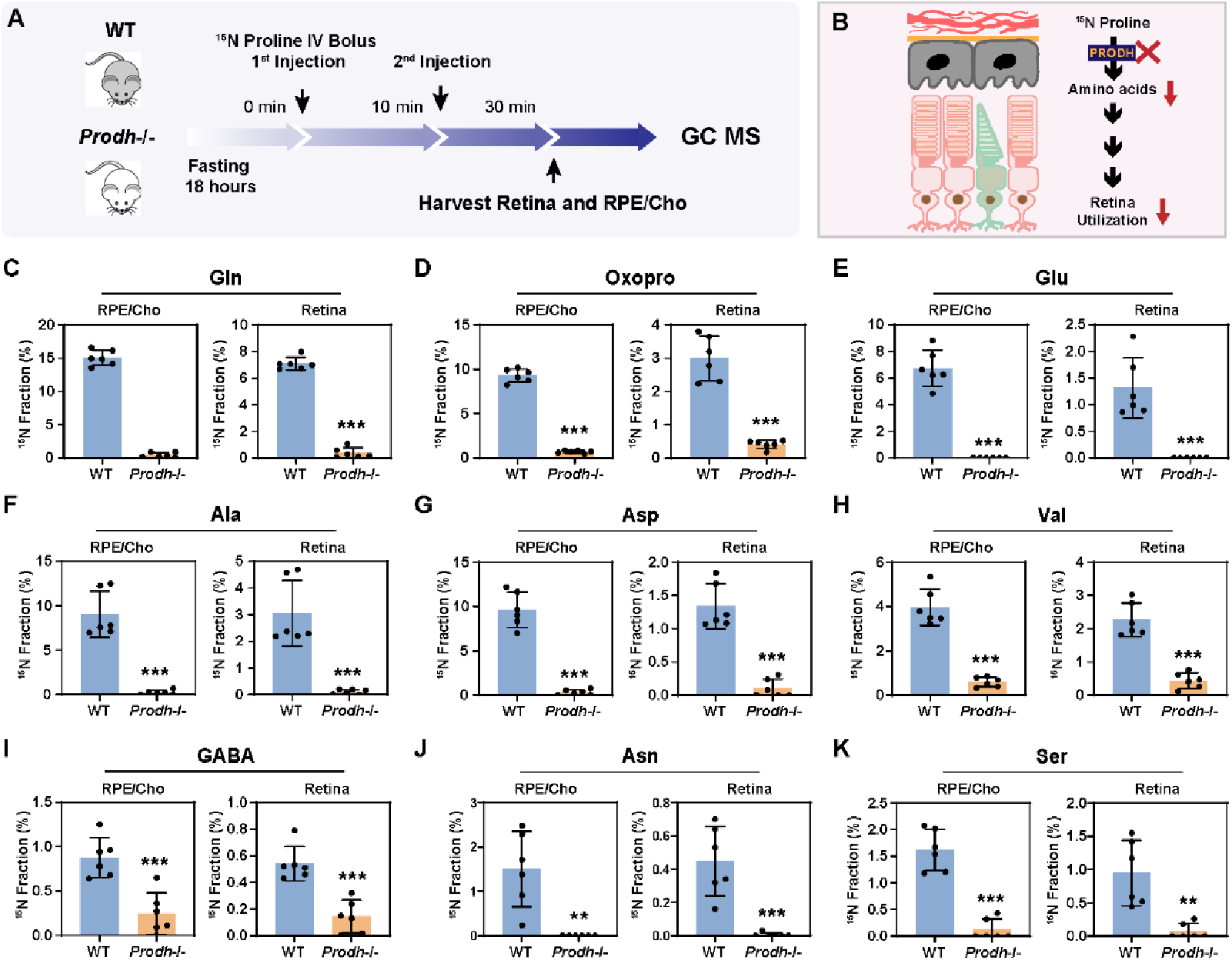
Depletion of PRODH blocks the synthesis of amino acids from proline in the RPE and its export into the retina. (A) A schematic of *in vivo* tracing of ^15^N proline. Mice were fasted for 18 hours and received two injections of ^15^N proline (150 mg/kg) at 0 min and 10 min, respectively. Retina and RPE/Cho were harvested at 30min after the second injection for metabolite analysis with GC MS. (B) *Prodh* -/-blocks the synthesis of amino acids from ^15^N proline in the RPE and its export into the retina. (C-K) ^15^N fraction of metabolites derived from ^15^N proline in RPE/Cho and retina. N=6.

## Discussion

In this study, we found that proline is a primary source of nitrogen source for synthesizing 13 types of amino acids in human RPE. More importantly, these proline-derived amino acids can be exported from the RPE to be used by the neural retina. The retina itself is unable to directly catabolize proline (**Fig 8**). Our findings demonstrate that in addition to its classical function as a nutrient transported between the choroid and the outer retina, the RPE serves as a major nutrient-manufacturing center, analogous to the liver, that supports retinal metabolism.

**Fig. 8.**
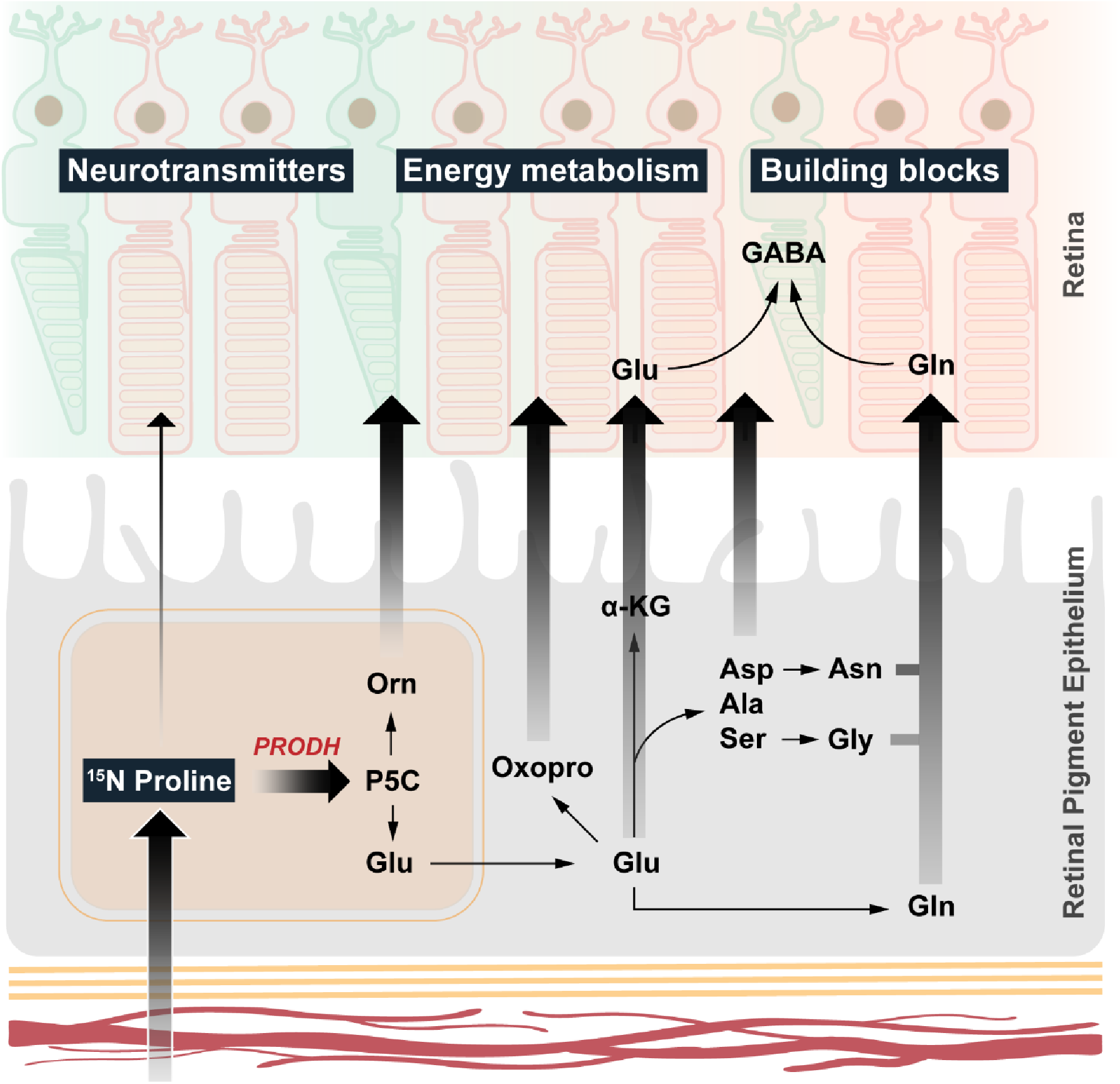
RPE synthesizes and export amino acids to the retina using nitrogen from proline. RPE uptakes and catabolizes proline through PRODH to produce P5C, which can be converted into ornithine, glutamate and multiple amino acids. All these amino acids can be exported to be utilized by the neural retina.

Photoreceptors in the outer retina are among the most energy-demanding cells in the human body, and their metabolic needs depend on their neighboring RPE. Due to the lack of a direct blood supply, photoreceptors must take up nutrients including glucose, amino acids, fatty acids, vitamins, and minerals from the choriocapillaris through the RPE. Deletion of the glucose transporter in the RPE is sufficient to cause photoreceptor degeneration. Besides directly transporting nutrients to the outer retina, the RPE also serves as a nutrient reservoir and manufacturer to support retinal metabolism. Transcriptomics data showed genes in glucose storage, proline metabolism and serine metabolism are highly upregulated in human RPE (6). Cultured RPE can store glucose as glycogen and oxidize glucose as mitochondrial intermediates to export them to the retinal side (5, 12). RPE can recycle photoreceptor-derived fatty acids and succinate into ketone bodies and malate to support the metabolic needs of the retina. Similar to its catabolism of carbon skeletons into mitochondrial intermediates (5), proline catabolizes its nitrogen into multiple amino acids in the RPE to export them for use by the retina. A common theme of these findings supports the idea that active RPE metabolism is essential for producing nutrients necessary to support the metabolically demanding neural retina.

Most of the enzymes that contribute to proline nitrogen metabolism are located in the mitochondria including PRODH, P5C dehydrogenase, OAT, and mitochondrial isoforms of transaminases. Inhibition of mitochondrial respiration can block proline catabolism and the function of nutrient release in RPE (13), suggesting that the supply of amino acids from proline catabolism to the retina requires healthy RPE mitochondrial metabolism. Mitochondrial metabolic defects in the RPE are known to cause RPE dysfunction including de-differentiation and lipid deposit formation, resulting in subsequent retinal degeneration and AMD in animal models and human donors (1, 14, 15). However, the mechanisms by which RPE mitochondrial dysfunction leads to photoreceptor degeneration remain unclear. Interestingly, de-differentiated RPE loses the ability to utilize proline (7) and the expression of transcripts in proline transport and catabolism are significantly reduced in de-differentiated RPE (16). We speculate that one consequence of mitochondrial defects in the RPE may be diminished nutrient production including proline catabolism, so that fewer amino acids and other nutrients are exported. That can induce metabolic stress for photoreceptors and cause degeneration.

Photoreceptors have a high metabolic need for amino acids. Every day, ∼10% of the photoreceptor outer segment material is shed and resynthesized. Rhodopsin makes up more than 90% of outer segment proteins (17) and human rhodopsin protein consists of 9.2% alanine, 23.8% BCAAs and 11.2% serine and glycine (18). Photoreceptors have extremely high energy expenditure with each photoreceptor consuming 1-1.5×10^8^ ATP/second (19). To achieve phototransduction, photoreceptors require the availability of a large amount of cGMP from GTP. The synthesis of ATP, GTP and cGMP requires nitrogen from aspartate, glutamine and glycine. Retinal cells including photoreceptors and glial cells need to maintain high concentrations of glutamate, aspartate, GABA, glutamine and glutathione for neurotransmission, the transfer of reducing equivalent and anti-oxidative stress (20-22). By unbiasedly analyzing the nutrient consumption of retinas from media that is nutrient-rich, we reported that glutamate and aspartate are specifically consumed by human retinas, mouse retina and mature human retinal organoid but not the immature organoid (6). This is consistent with our findings in this study that proline provides the nitrogen source for RPE synthesis of multiple amino acids that are utilized by the neural retina.

Why does the retina obtain nitrogen from proline metabolism in the RPE? Glutamine is abundant in the plasma, and it is a common nitrogen provider for proliferative tissues (23-25). Both RPE and the neural retina could readily utilize glutamine to produce glutamate, aspartate, and other amino acids (26). However, glutamine catabolism can generate free ammonia, which can be toxic to retinal neurons. Unlike the liver, the retina may not be able to condense the free ammonia into glutamate as glutamate dehydrogenase activity is almost absent in the retina (26). The removal of free ammonia in the retina relies on the synthesis of glutamine and asparagine (26). Alanine and BCAAs are also common nitrogen donors through alanine transaminase and branch-chain amino acid transaminase. However, we reported that the neural retina actively takes up alanine and BCAAs but cannot catabolize them to transfer their nitrogen groups to glutamate or other amino acids (26). RPE can metabolize alanine and BCAAs, and it remains to be determined how they contribute to nitrogen metabolism in the retina. Compared to other amino acids, proline has high solubility and its catabolism through PRODH can direct the transfer of electrons to FAD (9). The RPE is known to have abundant riboflavin and FAD for FAD-dependent metabolism, including succinate and fatty acid oxidation (27-29). Compared to glutamine, proline catabolism is more energy efficient as it generates FADH_2_ and NADH_2_ without producing free ammonia. Dietary proline supplementation in young pigs increases protein synthesis in a dose-dependent manner, while decreasing concentrations of urea in plasma (30). We found proline can be a nitrogen for 13 types of amino acids, covering 50% of proteogenic amino acids. Our findings suggest that proline can be an important nitrogen donor in retinal protein synthesis and metabolism.

Compared to human RPE cells, mouse RPE produces fewer ^15^N-labeled amino acids. This difference could be caused by several possibilities. The first is the short study durations in culture or *in vivo* delivery. In human RPE culture, ^15^N proline is incubated for 24-48 hours, which provides enough time for the equilibrium of reactions in proline metabolism. Because of limitations in culture techniques, we can only culture the RPE/Cho explants for 2 hours. The importance of reaction duration is more evident *in vivo*, in which double ^15^N proline injections increased 4 more labeled amino acids than from a single injection. Improvement of techniques in long-term explant culture and *in vivo* tracer perfusion should help address this limitation in the future. Secondly, there are differences in culture media compared to plasma. Plasma has a more complex nutrient composition and nitrogen sources than cell culture media. For example, ornithine is not typically added in culture media but is present at ∼100 μM in mouse plasma (31). In addition, there are differences in RPE composition and amounts of sample. Because of limitations in current techniques, it is almost impossible to isolate pure mouse RPE cells without affecting their metabolism. We have to use the RPE/Cho complex, which contains more extracellular matrix and capillaries than the pure human RPE culture. The RPE sample amount (∼54,000 RPE cells) from one mouse eye (32) is less than half of the number of human RPE cells in the culture. Finally, nitrogen metabolism between human and mouse RPE may be different intrinsically, as it has been shown that human and mouse RPE have differences in gene expression profiles (33), cell density, cell distribution (34) and susceptibility to diseases (35).

Among proline nitrogen-derived amino acids, GABA is the only amino acid with a decreased concentration decreased in the co-cultured retinas but increased in the co-cultured RPE (**Fig 3C**). These results suggest a GABA efflux from the retina to the RPE. Consistently, in the RPE from ^15^N proline-injected mice, the percentage of ^15^N labeling drops at 60 min for all the amino acids except for GABA (**Fig 5**). However, the concentration of retinal GABA remains constant or slightly increased after injection. One possibility is that there is a rapid rate of GABA synthesis and recycling between the RPE and retina *in vivo*. GABA can also be produced through its transaminase from succinic semialdehyde, which is not in the culture media. GABA is located mostly in ganglion cells, amacrine cells, horizontal cells, and Müller glial cells (36, 37). Ganglion cells and other inner neurons are easily stressed and damaged after removing the optic nerve in the retinal explant culture. Another possibility for the decrease of GABA content *in vitro* may be caused by the stimulation and damage of these inner retinal neurons after isolation.

In conclusion, we demonstrate that the neural retina needs nitrogen sources to maintain its amino acid pools, but it cannot directly catabolize proline. The RPE actively catabolizes proline to synthesize multiple amino acids that are exported to support retinal metabolism. These findings highlight the importance of RPE metabolism in supporting nitrogen utilization in the neural retina, providing insight into the mechanism of RPE-initiated retinal degenerative diseases such as AMD.

## Abbreviation

RPE: (retinal pigment epithelium)
TCA: (tricarboxylic acid)
GC MS: (gas chromatography mass spectrometry)
Cho: (choroid)
Pro: (Proline)
Gln: (Glutamine)
Asp: (Aspartate)
Ala: (Alanine)
Oxopro: (Oxoproline)
Glu: (Glutamate)
Val: (Valine)
Ileu: (Isoleucine)
Leu: (Leucine)
Orn: (Ornithine)
Asn: (Asparagine)
GABA: (γ-Aminobutyric acid)
Ser: (Serine)
Gly: (Glycine)
PRODH: (proline dehydrogenase)

## Experimental Procedures

### Reagents

All the reagents and resources were detailed in the Key resources table (**Table S1**).

### Animals

Both genders of ∼2-month-old WT mice (C57 B6/J background, Stock #:000664) and *Prodh*-/-mice (C57BL/6J and 129P1/ReJ background, Stock #001137) were purchased from the Jackson lab. Mouse experiments were performed in accordance with the National Institutes of Health guidelines and the protocol was approved by the Institutional Animal Care and Use Committee of West Virginia University. For tracer study *in vivo*, mice were deprived of food for 18 hours and received bolus or double intravenous injections of ^15^N proline retro-orbitally at 150 mg/kg under anesthesia with isoflurane. Mouse retina and RPE/Cho were quickly harvested and snap-frozen in liquid nitrogen and stored at -80 °C before use.

### Human RPE cell culture

The use of RPE cells was reviewed by the University of Washington Institutional Review Board and experimental protocols follow ethical guidelines from the Declaration of Helsinki.

Human RPE cells were cultured as previously described (5, 7). Briefly, human fetal RPE cells were passaged using 0.25% trypsin and plated in 5% fetal bovine serum (v/v) in MEM-alpha supplemented with N1, NEAA, 1% pen/strep (v/v) and a mix of taurine, hydrocortisone and triiodothyronine (THT) with rock inhibitor for the first week. The media is then changed to the same recipe with reduced 1% (v/v) FBS in MEM-alpha media for maintenance. Human fetal RPE was grown in 12 well plates on Martigel for 20 weeks for experiment. Upon experiment, media was changed to clear DMEM (no glucose, sodium pyruvate, glutamine or phenol red) and then supplemented with 5.5mM glucose and 1mM unlabeled or 1mM ^15^N-proline with 1% FBS and Pen/strep (v/v). Media were collected at 0, 24 and 48 hrs of culturing for metabolomic analysis. Human RPE cells were washed with 154mM NaCl solution and collected on dry ice after 48 hrs of culturing for the same purpose.

### Retina and RPE/Cho explant culture

Mouse retina and RPE/Cho were quickly isolated in 200 μL of cold Hank’s Balanced Salt Solution (HBSS) as previously described (7, 26). The retina and RPE/Cho were transferred into 200 μL of pre-incubated KRB (7) with ^15^N proline in the presence of 5 mM glucose, followed by a 2-hour incubation at 37 °C in a CO_2_ incubator. The tissues were collected and snap-frozen in liquid nitrogen and stored at -80 °C before use.

### RPE and retina co-culture

Human RPE (hfRPE) was grown for 12-16 weeks in 12-well plates coated with matrigel before being used for co-culture experiments. RPE was equilibrated in unlabeled proline media for approximately 1 hr before starting the co-culture. Murine retinas were harvested from 6 to 8-week-old C57Bl/6J mice and allowed to equilibrate in unlabeled proline media for about 1hr. After equilibration, the cells alone or co-cultured with the retina were switched to ^15^N proline media and incubated for 6 hrs. After 6 hrs, media was harvested to a separate tube from the tissue. The retinas were quickly removed from hfRPE and placed into a cold 154mM NaCl solution, spun down, liquid removed and snap frozen. RPE cells were washed with the same saline solution, and then harvested on ice with cold 80% methanol and stored frozen at -80 until processing.

### Metabolite analysis with GC MS

Metabolites from tissue, cells and media were extracted in 80% cold methanol as described previously (5). Metabolite extracts were transferred into glass inserts containing internal standard (Myristic acid d27 in 1 mg/ml) and dried in a speedvac. Dried samples were derivatized by methoxyamine and N-tertbutyldimethylsilyl-N-methyl trifluoro-acetamide and analyzed by the Agilent 7890B/5977B GC MS system with DB-5MS column (30 m × 0.25 mm × 0.25 μm film) as we described in detail before (5, 26). Mass spectra were collected from 80–600 m/z under SIM mode. Table S2 lists the detailed parameters including monitored ions for the measured metabolites. The data was analyzed by Agilent MassHunter Quantitative Analysis Software by extracting ion abundance for each monitored ion in **Table S2**. Raw total ion abundance for isotopologues, the purity of the tracers and derivatization reagents were imported in ISOCOR software (https://isocor.readthedocs.io/en/latest/) to calculate the percentage of labeling by correcting natural abundance and tracer purity (26). The percentage of labeled ions or enrichment was presented as ^15^N of the total. The total abundance was the total ion abundance from all isotopologues.

### Western blot

Western blot was performed as previously described (38). Briefly, proteins were extracted from tissues using RIPA buffer and determined for protein concentration by the BCA protein assay kit. Protein samples (20μg) were boiled and loaded onto SDS-PAGE gels. The gels were transferred to nitrocellulose membranes and blocked with 5% non-fat milk in 1X Tris-Buffered Saline containing 0.1% Tween® 20 (TBST). The membranes were incubated with primary antibodies (See details in **Table S1**) at 4°C overnight. After washing with 1×PBS containing 0.1% Tween-20 (PBST), the membranes were incubated with a rabbit or mouse secondary antibody conjugated with horseradish peroxidase (1:2000) for 1 hour, followed by three washes with PBST. A chemiluminescence reagent kit was used to visualize protein bands with horseradish peroxidase secondary antibodies.

### Statistics

The significance of differences between means was determined by unpaired two-tailed T-tests or One-Way ANOVA with Bonferroni post hoc test using GraphPad Prism 9.0 (GraphPad Software, Inc). Only P < 0.05 was considered to be significant. All data are presented as the mean ± SD.

## Supporting Information

This article contains supporting information.

## Author Contributions

Conceptualization, J.C. and J.D; Investigation, S.Z., R.X., A.L.E., Y.W., R.M., J.H., J.C., J.D.; Writing, S.Z., R.X., A.L.E, J.B.H., J. C.., and J.D.; Funding Acquisition J.C., and J.D; Supervision, J.C. and J.D.

## Acknowledgments

This work was supported by National Eye Institute Grant EY034364 (JC, JD), EY031324 (JD), EY032462(JD), Bright Focus Foundation M2020141(JD), the Retina Research Foundation (JD), National Institute of General Medical Sciences P20GM144230 (Visual Sciences Center of Biomedical Research Excellence), and funds for Core facilities P20 GM103434 (WV INBRE grant).

## Ethical approval

The mouse experiments were performed in accordance with the National Institutes of Health guidelines and the protocol approved by the Institutional Animal Care and Use Committee of West Virginia University.

The use of human RPE cells were reviewed by the University of Washington Institutional Review Board and experimental protocols follow ethical guidelines from the Declaration of Helsinki.

## Informed consent

The authors declare no patients involved in this study and no informed consent was made.

## Data Availability Statement

GC MS data generated in this study were provided in the supplemental material.

## References

1. Ferrington, D. A., Fisher, C. R., andKowluru, R. A. (2020) Mitochondrial Defects Drive Degenerative Retinal Diseases Trends Mol Med 26, 105–118 10.1016/j.molmed.2019.10.008

2. Lin, H., Xu, H., Liang, F. Q., Liang, H., Gupta, P., Havey, A. N. et al. (2011) Mitochondrial DNA damage and repair in RPE associated with aging and age-related macular degeneration Invest Ophthalmol Vis Sci 52, 3521–3529 10.1167/iovs.10-6163

3. Hurley, J. B. (2021) Retina Metabolism and Metabolism in the Pigmented Epithelium: A Busy Intersection Annu Rev Vis Sci 7, 665–692 10.1146/annurev-vision-100419-115156

4. Kanow, M. A., Giarmarco, M. M., Jankowski, C. S., Tsantilas, K., Engel, A. L., Du, J. et al. (2017) Biochemical adaptations of the retina and retinal pigment epithelium support a metabolic ecosystem in the vertebrate eye Elife 6, 10.7554/eLife.28899

5. Chao, J. R., Knight, K., Engel, A. L., Jankowski, C., Wang, Y., Manson, M. A. et al. (2017) Human retinal pigment epithelial cells prefer proline as a nutrient and transport metabolic intermediates to the retinal side J Biol Chem 292, 12895–12905 10.1074/jbc.M117.788422

6. Li, B., Zhang, T., Liu, W., Wang, Y., Xu, R., Zeng, S. et al. (2020) Metabolic Features of Mouse and Human Retinas: Rods versus Cones, Macula versus Periphery, Retina versus RPE iScience 23, 101672 10.1016/j.isci.2020.101672

7. Yam, M., Engel, A. L., Wang, Y., Zhu, S., Hauer, A., Zhang, R. et al. (2019) Proline mediates metabolic communication between retinal pigment epithelial cells and the retina J Biol Chem 294, 10278–10289 10.1074/jbc.RA119.007983

8. Kennaway, N. G., Stankova, L., Wirtz, M. K., andWeleber, R. G. (1989) Gyrate atrophy of the choroid and retina: characterization of mutant ornithine aminotransferase and mechanism of response to vitamin B6 Am J Hum Genet 44, 344–352, https://www.ncbi.nlm.nih.gov/pubmed/2916580

9. Du, J., Zhu, S., Lim, R. R., andChao, J. R. (2021) Proline metabolism and transport in retinal health and disease Amino Acids 53, 1789–1806 10.1007/s00726-021-02981-1

10. Gogos, J. A., Santha, M., Takacs, Z., Beck, K. D., Luine, V., Lucas, L. R. et al. (1999) The gene encoding proline dehydrogenase modulates sensorimotor gating in mice Nat Genet 21, 434–439 10.1038/7777

11. Blake, R. L., andRussell, E. S. (1972) Hyperprolinemia and prolinuria in a new inbred strain of mice, PRO-Re Science 176, 809–811 10.1126/science.176.4036.809

12. Senanayake, P., Calabro, A., Hu, J. G., Bonilha, V. L., Darr, A., Bok, D. et al. (2006) Glucose utilization by the retinal pigment epithelium: evidence for rapid uptake and storage in glycogen, followed by glycogen utilization Exp Eye Res 83, 235–246 10.1016/j.exer.2005.10.034

13. Zhang, R., Engel, A. L., Wang, Y., Li, B., Shen, W., Gillies, M. C. et al. (2021) Inhibition of Mitochondrial Respiration Impairs Nutrient Consumption and Metabolite Transport in Human Retinal Pigment Epithelium J Proteome Res 20, 909–922 10.1021/acs.jproteome.0c00690

14. Zhao, C., Yasumura, D., Li, X., Matthes, M., Lloyd, M., Nielsen, G. et al. (2011) mTOR-mediated dedifferentiation of the retinal pigment epithelium initiates photoreceptor degeneration in mice J Clin Invest 121, 369–383 10.1172/JCI44303

15. Felszeghy, S., Viiri, J., Paterno, J. J., Hyttinen, J. M. T., Koskela, A., Chen, M. et al. (2019) Loss of NRF-2 and PGC-1alpha genes leads to retinal pigment epithelium damage resembling dry age-related macular degeneration Redox Biol 20, 1–12 10.1016/j.redox.2018.09.011

16. Boles, N. C., Fernandes, M., Swigut, T., Srinivasan, R., Schiff, L., Rada-Iglesias, A. et al. (2020) Epigenomic and Transcriptomic Changes During Human RPE EMT in a Stem Cell Model of Epiretinal Membrane Pathogenesis and Prevention by Nicotinamide Stem Cell Reports 14, 631–647 10.1016/j.stemcr.2020.03.009

17. Filipek, S., Stenkamp, R. E., Teller, D. C., andPalczewski, K. (2003) G protein-coupled receptor rhodopsin: a prospectus Annu Rev Physiol 65, 851–879 10.1146/annurev.physiol.65.092101.142611

18. Nathans, J., andHogness, D. S. (1984) Isolation and nucleotide sequence of the gene encoding human rhodopsin Proc Natl Acad Sci U S A 81, 4851–4855 10.1073/pnas.81.15.4851

19. Ingram, N. T., Fain, G. L., andSampath, A. P. (2020) Elevated energy requirement of cone photoreceptors Proc Natl Acad Sci U S A 117, 19599–19603 10.1073/pnas.2001776117

20. Winkler, B. S., andGiblin, F. J. (1983) Glutathione oxidation in retina: effects on biochemical and electrical activities Exp Eye Res 36, 287–297 10.1016/0014-4835(83)90013-1

21. Kalloniatis, M., Marc, R. E., andMurry, R. F. (1996) Amino acid signatures in the primate retina J Neurosci 16, 6807–6829 10.1523/JNEUROSCI.16-21-06807.1996

22. Heinamaki, A. A., Muhonen, A. S., andPiha, R. S. (1986) Taurine and other free amino acids in the retina, vitreous, lens, iris-ciliary body, and cornea of the rat eye Neurochem Res 11, 535–542 10.1007/BF00965323

23. Cruzat, V., Macedo Rogero, M., Noel Keane, K., Curi, R., andNewsholme, P. (2018) Glutamine: Metabolism and Immune Function, Supplementation and Clinical Translation Nutrients 10, 10.3390/nu10111564

24. Casado, J., Felipe, A., Pastor-Anglada, M., andRemesar, X. (1988) Glutamine as a major nitrogen carrier to the liver in suckling rat pups Biochem J 256, 377–381 10.1042/bj2560377

25. Yoo, H. C., Yu, Y. C., Sung, Y., andHan, J. M. (2020) Glutamine reliance in cell metabolism Exp Mol Med 52, 1496–1516 10.1038/s12276-020-00504-8

26. Xu, R., Ritz, B. K., Wang, Y., Huang, J., Zhao, C., Gong, K. et al. (2020) The retina and retinal pigment epithelium differ in nitrogen metabolism and are metabolically connected J Biol Chem 295, 2324–2335 10.1074/jbc.RA119.011727

27. Wang, Y., Grenell, A., Zhong, F., Yam, M., Hauer, A., Gregor, E. et al. (2018) Metabolic signature of the aging eye in mice Neurobiol Aging 71, 223–233 10.1016/j.neurobiolaging.2018.07.024

28. Hass, D. T., Bisbach, C. M., Robbings, B. M., Sadilek, M., Sweet, I. R., andHurley, J. B. (2022) Succinate metabolism in the retinal pigment epithelium uncouples respiration from ATP synthesis Cell Rep 39, 110917 10.1016/j.celrep.2022.110917

29. Reyes-Reveles, J., Dhingra, A., Alexander, D., Bragin, A., Philp, N. J., andBoesze-Battaglia, K. (2017) Phagocytosis-dependent ketogenesis in retinal pigment epithelium J Biol Chem 292, 8038–8047 10.1074/jbc.M116.770784

30. Wu, G., Bazer, F. W., Burghardt, R. C., Johnson, G. A., Kim, S. W., Knabe, D. A. et al. (2011) Proline and hydroxyproline metabolism: implications for animal and human nutrition Amino Acids 40, 1053–1063 10.1007/s00726-010-0715-z

31. Wang, T., Steel, G., Milam, A. H., andValle, D. (2000) Correction of ornithine accumulation prevents retinal degeneration in a mouse model of gyrate atrophy of the choroid and retina Proc Natl Acad Sci U S A 97, 1224–1229 10.1073/pnas.97.3.1224

32. Bodenstein, L., andSidman, R. L. (1987) Growth and development of the mouse retinal pigment epithelium. I. Cell and tissue morphometrics and topography of mitotic activity Dev Biol 121, 192–204 10.1016/0012-1606(87)90152-7

33. Bennis, A., Gorgels, T. G., Ten Brink, J. B., van der Spek, P. J., Bossers, K., Heine, V. M. et al. (2015) Comparison of Mouse and Human Retinal Pigment Epithelium Gene Expression Profiles: Potential Implications for Age-Related Macular Degeneration PLoS One 10, e0141597 10.1371/journal.pone.0141597

34. Zhang, J., Xu, G., Zhang, L., Gu, L., Xu, H., Lu, L. et al. (2012) A modified histoimmunochemistry-assisted method for in situ RPE evaluation Front Biosci (Elite Ed) 4, 1571–1581 10.2741/481

35. Zhang, Y., Stanton, J. B., Wu, J., Yu, K., Hartzell, H. C., Peachey, N. S. et al. (2010) Suppression of Ca2+ signaling in a mouse model of Best disease Hum Mol Genet 19, 1108–1118 10.1093/hmg/ddp583

36. Lam, D. M. (1972) The biosynthesis and content of gamma-aminobutyric acid in the goldifsh retina J Cell Biol 54, 225–231 10.1083/jcb.54.2.225

37. Bolz, J., Frumkes, T., Voigt, T., andWassle, H. (1985) Action and localization of gamma-aminobutyric acid in the cat retina J Physiol 362, 369–393 10.1113/jphysiol.1985.sp015684

38. Zhu, S., Huang, J., Xu, R., Wang, Y., Wan, Y., McNeel, R. et al. (2022) Isocitrate dehydrogenase 3b is required for spermiogenesis but dispensable for retinal viability J Biol Chem 298, 102387 10.1016/j.jbc.2022.102387

