## Supplementary information for "Proline provides a nitrogen source in the retinal pigment epithelium to synthesize and export amino acids for the neural retina"

This PDF file includes:

Supplementary Figures 1-5

Supplementary Table 1-2

**Supplementary Figures**

**Supplementary Figure 1**

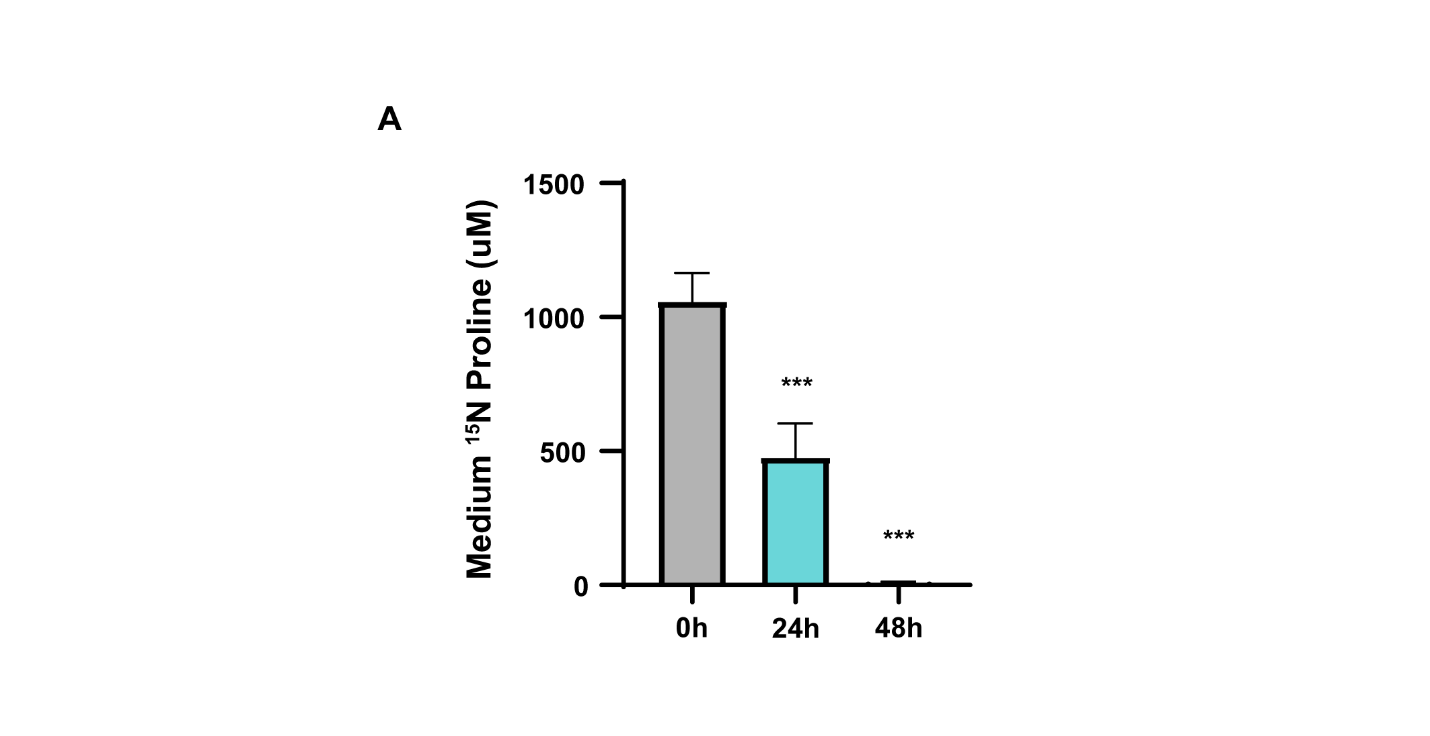

**Figure S1. RPE quickly consumes proline from the media.** Human RPE cells were incubated with or without 1 mM ^15^N proline in DMEM supplemented with 5.5 mM glucose and 1% FBS. The spent media were collected at 0h, 24h and 48h for metabolite analysis with GC MS. (A) Proline level in the medium after different time points of culturing. ***P<0.001, N=3.

**Supplementary Figure 2**

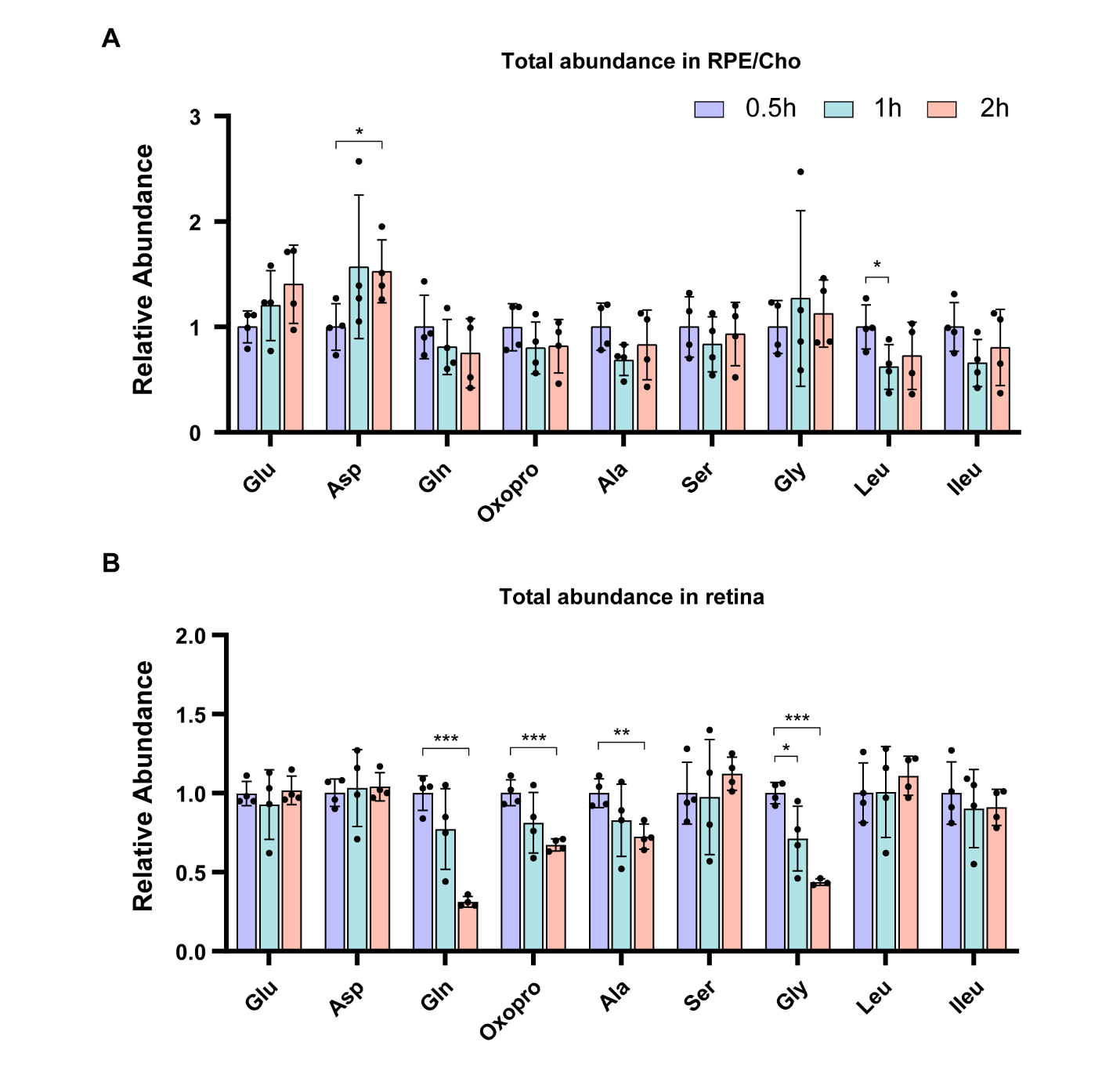

**Figure S2. Total abundance of amino acids in mouse RPE/Cho and retina.** Tissues were incubated with 1 mM ^15^N proline in KRB with 5.5 mM glucose for 0.5h, 1h and 2h. Tissues and media were collected at these time points for metabolite analysis with GC MS. (A-B) Total abundance of all isotopologue metabolites in mouse RPE/Cho and retina at different time points relative to 0.5h incubation. *P<0.05, **P<0.01, ***P<0.001, N=4. Glu (Glutamate), Asp (Aspartate), Gln (Glutamine), Oxopro (Oxoproline), Ala (Alanine), Ser (Serine), Gly (Glycine), Leu (Leucine), Ileu (Isoleucine).

**Supplementary Figure 3**
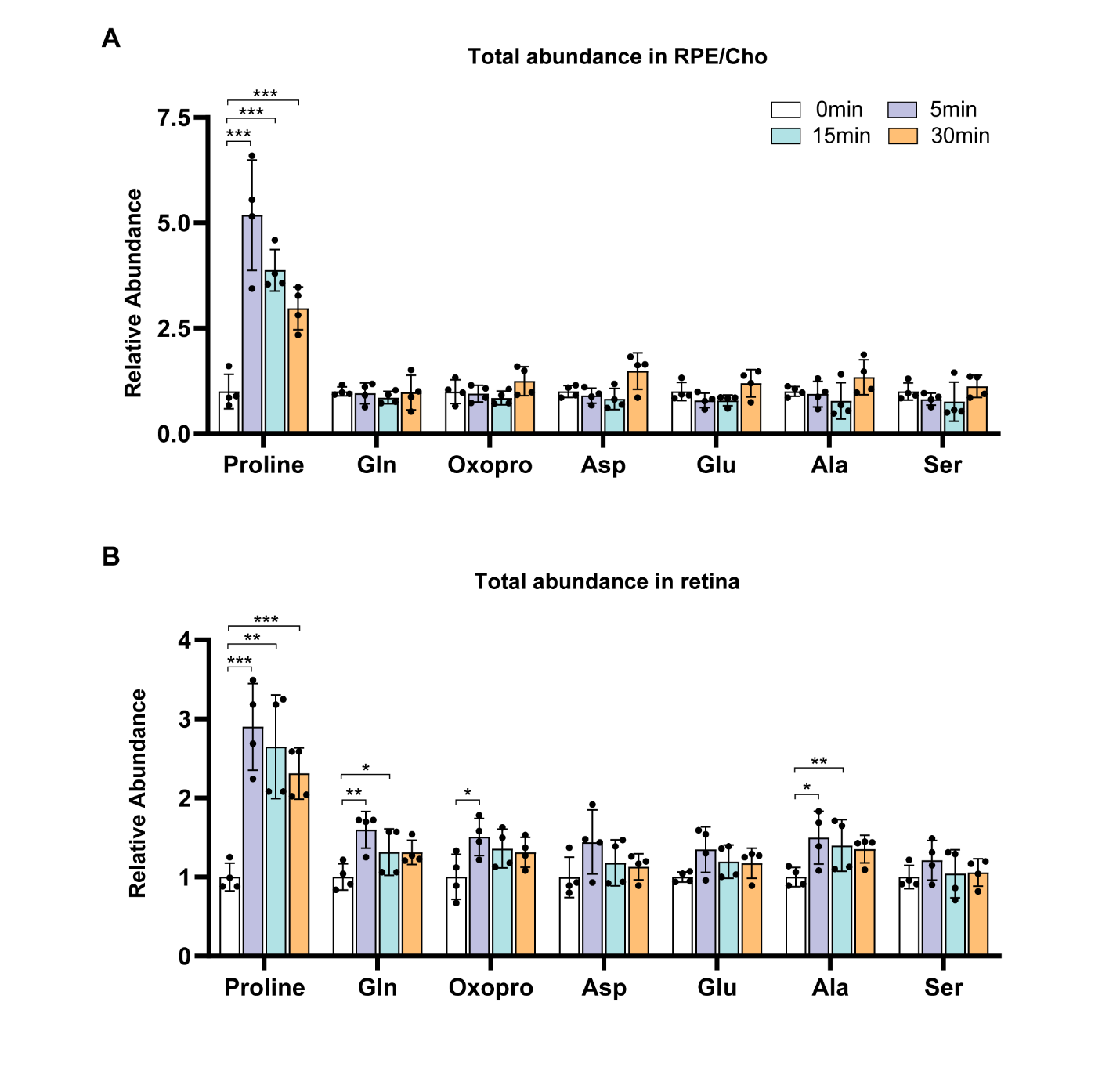
**Figure S3. Total abundance of amino acids in mouse RPE/Cho and retina after injection of ^15^N proline.** WT mice were fasted for 18 hours and received a single retro-orbital injection of ^15^N proline (150 mg/kg) or PBS. Retina and RPE/Cho were harvested at 0min, 5min, 15min and 30min after injection for metabolite analysis with GC MS. (A-B) Total abundance of all isotopologue metabolites in mouse RPE/Cho and retina at different time points relative to 0 min after injection. *P<0.05, **P<0.01, ***P<0.001, N=4.

**Supplementary Figure 4**

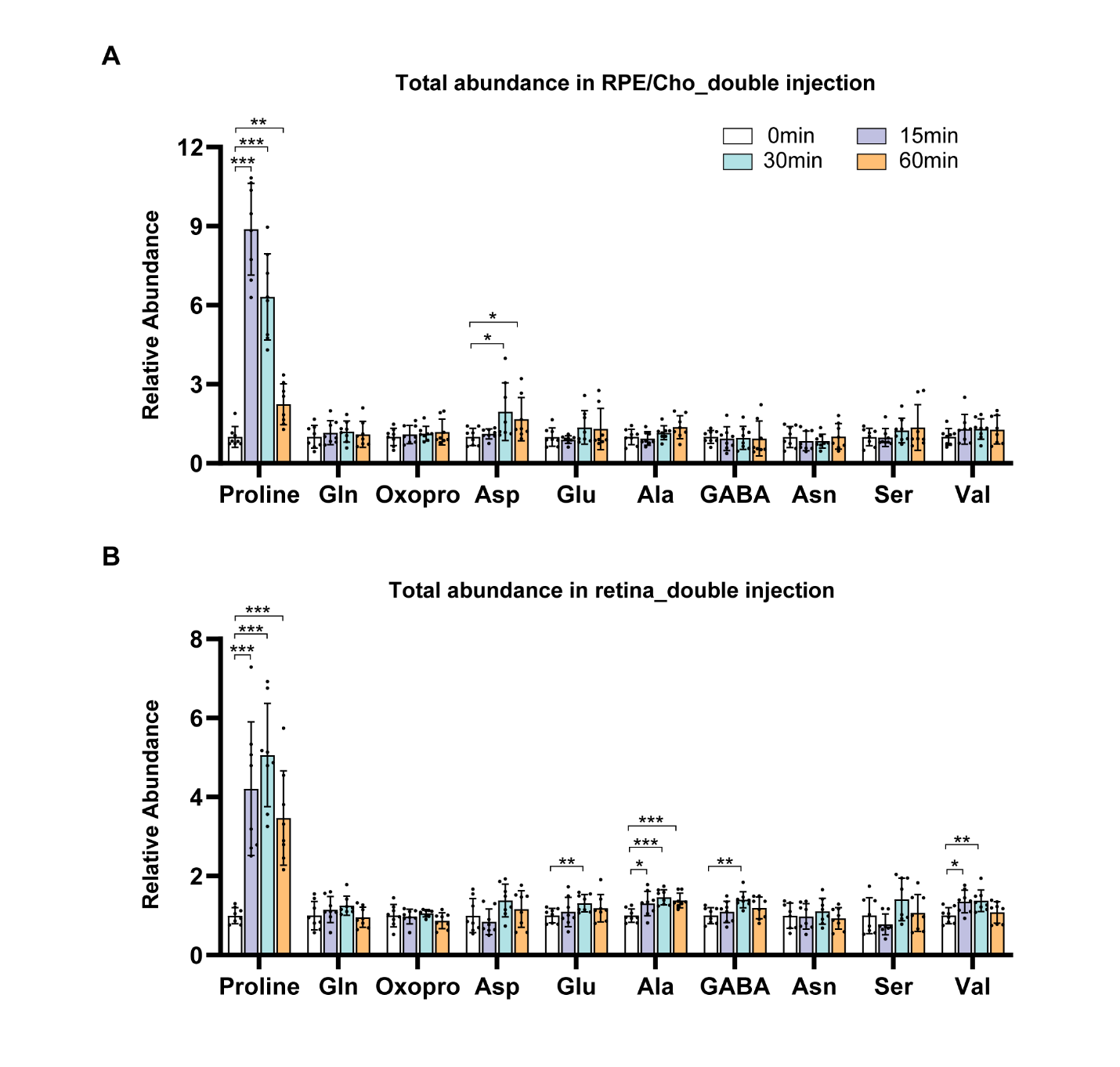
**Figure S4. Total abundance of amino acids in mouse RPE/Cho and retina after injection of ^15^N proline.** WT mice were fasted for 18 hours and received two injections of ^15^N proline (150 mg/kg) or PBS at 0 min and 10 min, respectively. Retina and RPE/Cho were harvested at 0min, 15min, 30min and 60min after the second injection for metabolite analysis with GC MS. (A-B) Total abundance of all isotopologue metabolites in mouse RPE/Cho and retina at different time points relative to 0 min after injection. *P<0.05, **P<0.01, ***P<0.001, N=8.

**Supplementary Figure 5**

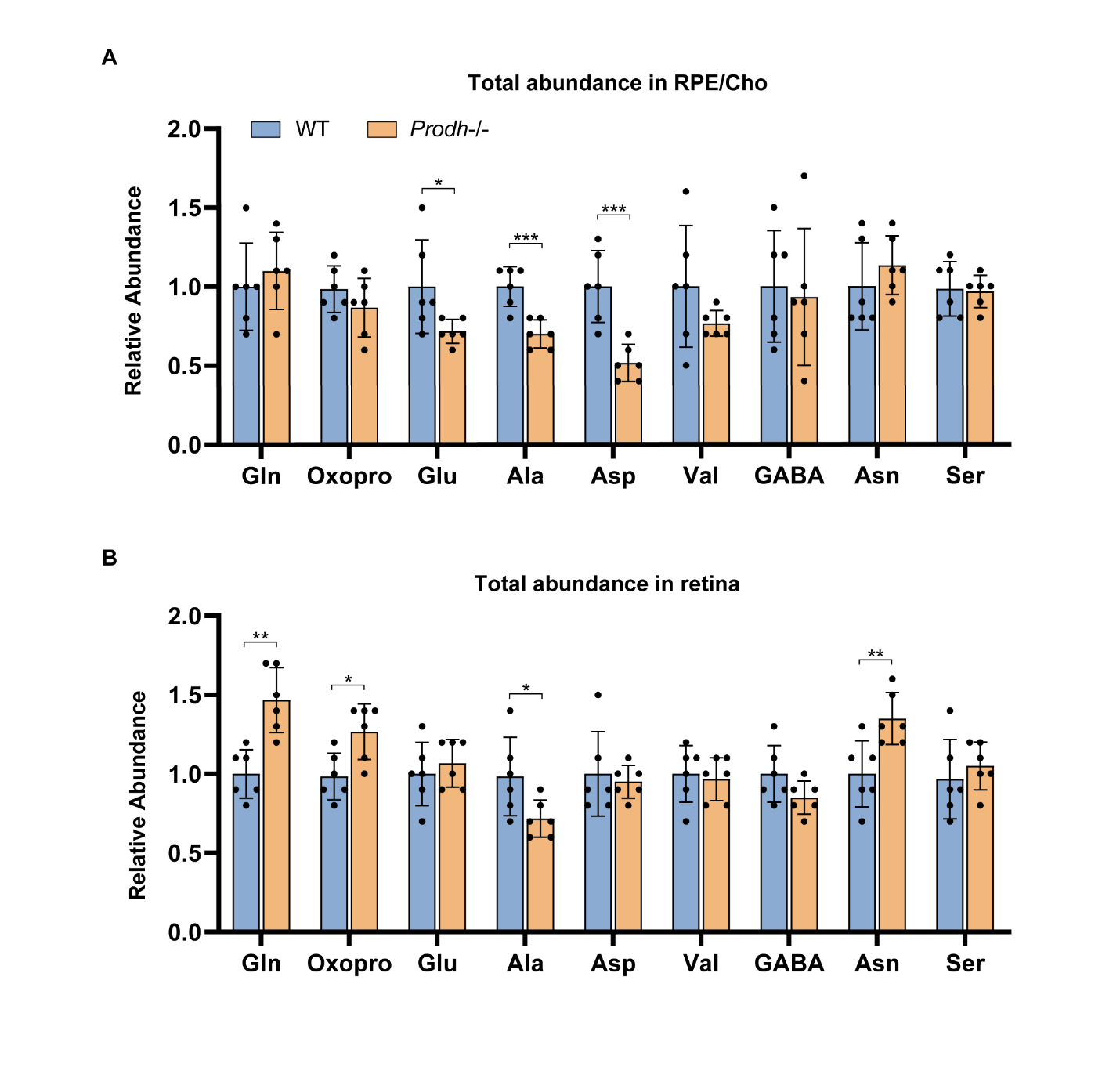
**Figure S5. Total abundance of amino acids in RPE/Cho and retina from WT and Prodh -/- mice after double injection of ^15^N proline.** Mice were fasted for 18 hours and received two injections of ^15^N proline (150 mg/kg) or PBS at 0 min and 10 min, respectively. Retina and RPE/Cho were harvested at 30 min after the second injection for metabolite analysis with GC MS. (A-B) Total abundance of all isotopologue metabolites in *Prodh*-/- mouse RPE/Cho and retina relative to WT mouse. *P<0.05, **P<0.01, ***P<0.001, N=6.

**Table S1. Key resources**

| **Reagents** | **Catalog** | **Company** | **Location** |
| --- | --- | --- | --- |
| ^15^N Proline | 608998 | Sigma-Alderich | St. Louis, MO USA |
| Minimum Essential Medium Eagle (MEM alpha) | M4526 | Sigma-Aldrich | St. Louis, MO USA |
| N1 Medium Supplement (100×) | N6530-5ML | Sigma-Aldrich | St. Louis, MO USA |
| MEM Non-Essential Amino Acids Solution (100X) (NEAA) | 11140050 | Thermo scientific | Rockford, IL USA |
| Fetal Bovine Serum (FBS) | S11550 | Atlanta Biologicals | Flowery Branch, GA USA |
| Taurine | T0625-10G | Sigma-Aldrich | St. Louis, MO USA |
| 3, 3', 5-Triiodo-L-Thyronine | T6397-100MG | Sigma-Aldrich | St. Louis, MO USA |
| Y-27632 dihydrochloride (rock inhibitor) | 1254 | Tocris/R&D Systems | Minneapolis, MN USA |
| DMEM, no glucose, no glutamine, no phenol red | A1443001 | Thermo Fisher Scientific | Rockford IL USA |
| Penicillin-Streptomycin (5,000 U/mL) | 15070-063 | Thermo Fisher Scientific | Rockford, IL USA |
| RIPA Lysis and Extraction Buffer | 89900 | Thermo Fisher Scientific ic | Rockford, IL USA |
| Pierce Protease and Phosphatase Inhibitor Mini Tablets | A32959 | Thermo Fisher Scientific | Rockford, IL USA |
| Pierce™ BCA Protein Assay Kit | 23225 | Thermo Fisher Scientific | Rockford, IL USA |
| TGX Stain-Free™ FastCast™ Acrylamide Kit, 10% | 1610183 | Bio-Rad | Hercules, CA USA |
| 10x Tris/Glycine/SDS | 1610732 | Bio-Rad | Hercules, CA USA |
| 10x Tris/Glycine Buffer for Western Blots and Native Gels | 1610734 | Bio-Rad | Hercules, CA USA |
| Nitrocellulose membrane | 1620097 | Bio-Rad | Hercules, CA USA |
| Blotting-Grade Blocker (non-fat dry milk) | 1706404 | Bio-Rad | Hercules, CA USA |
| Immobilon Western Chemiluminescent HRP Substrate | WBKLS0500 | EMD Millipore Corporation | Burlington, MA USA |
| Phosphate buffer saline | P3813-10PAK | Sigma-Aldrich | St. Louis, MO USA |
| Methanol | 14262-1L | Honeywell | Morris Plains, NJ USA |
| Methoxyamine hydrochloride | 226904-1G | Sigma-Aldrich | St. Louis, MO USA |
| N-tert-butyldimethylsilyl-N-methyltrifluoroacetamide (TBDMS) | 190500 | Sigma-Aldrich | St. Louis, MO USA |
| **Antibodies&Host** | **Company** | **Catalog** | **Dilutions** |
| Anti-PRODH Antibody, rabbit | Sigma-Aldrich | 22980-I-AP | 1:1000 (WB) |
| Anti-β-Tubulin Antibody, mouse | Santa Cruz | sc-55529 | 1:1000 (WB) |
| Anti-GAPDH Antibody, mouse | Santa Cruz | sc-32233 | 1:1000 (WB) |
| Goat anti-Rabbit HRP, goat | Cell Signaling Technology | 7074S | 1:2000 (WB) |
| Goat anti-Mouse HRP, goat | Abcam | ab131368 | 1:2000 (WB) |

**WB: Western Blot**

**Table S2. Parameters for amino acids measured in GC MS.**

| **Metabolite** | **Monitored ion (m/z)** | **Platform** | **Polarity** |
| --- | --- | --- | --- |
| Alanine M0 | 260.2 | GCMS | + |
| Alanine M1 | 261.2 | GCMS | + |
| Asparagine M0 | 417.2 | GCMS | + |
| Asparagine M1 | 418.2 | GCMS | + |
| Asparagine M2 | 419.2 | GCMS | + |
| Aspartate M0 | 418.2 | GCMS | + |
| Aspartate M1 | 419.2 | GCMS | + |
| GABA M0 | 274.2 | GCMS | + |
| GABA M1 | 275.2 | GCMS | + |
| Glutamate M0 | 432.3 | GCMS | + |
| Glutamate M1 | 433.3 | GCMS | + |
| Glutamine M0 | 431.3 | GCMS | + |
| Glutamine M1 | 432.3 | GCMS | + |
| Glutamine M2 | 433.3 | GCMS | + |
| Glycine M0 | 246.1 | GCMS | + |
| Glycine M1 | 247.1 | GCMS | + |
| Isoleucine M0 | 200.2 | GCMS | + |
| Isoleucine M1 | 201.2 | GCMS | + |
| Leucine M0 | 200.2 | GCMS | + |
| Leucine M1 | 201.2 | GCMS | + |
| Proline M0 | 184.1 | GCMS | + |
| Proline M1 | 185.1 | GCMS | + |
| 5-Oxoproline M0 | 300.2 | GCMS | + |
| 5-Oxoproline M1 | 301.2 | GCMS | + |
| Serine M0 | 390.2 | GCMS | + |
| Serine M1 | 391.2 | GCMS | + |
| Valine M0 | 186.2 | GCMS | + |
| Valine M1 | 187.2 | GCMS | + |
| Ornithine M0 | 417.3 | GCMS | + |
| Ornithine M1 | 418.3 | GCMS | + |
| Ornithine M2 | 419.3 | GCMS | + |
